# Early life stress induced sex-specific changes in behavior is paralleled by altered locus coeruleus physiology in BALB/cJ mice

**DOI:** 10.1101/2024.08.06.606908

**Authors:** Brannan Savannah, Lauren Garbe, Ben D Richardson

**Affiliations:** Department of Pharmacology, Southern Illinois University – School of Medicine, Springfield, IL 62702

## Abstract

Adverse childhood experiences have been associated with many neurodevelopmental and affective disorders including attention deficit hyperactivity disorder and generalized anxiety disorder, with more exposures increasing negative risks. Sex and genetic background are biological variables involved in adverse psychiatric outcomes due to early life trauma. Females in general have an increased prevalence of stress-related psychopathologies beginning after adolescence, indicative of the adolescent time period being a female-specific sensitive period. To understand the underlying neuronal components responsible for this relationship between genetic background, sex, stress/trauma, and cognitive/affective behaviors, we assessed behavioral and neuronal changes in a novel animal model of early life stress exposure. Male and female BALB/cJ mice that express elevated basal anxiety-like behaviors and differences in monoamine associated genes, were exposed to an early life variable stress protocol that combined deprivation in early life with unpredictability in adolescence. Stress exposure produced hyperlocomotion and attention (5 choice serial reaction time task) in male and female mice along with female-specific increased anxiety-like behavior. These behavioral changes were paralleled by reduced excitability of locus coeruleus (LC) neurons, due to resting membrane potential hyperpolarization in males and a female specific increase in action potential delay time. These data describe a novel interaction between sex, genetic background, and early life stress that results in behavioral changes in clinically-relevant domains and potential underlying mechanistic lasting changes in physiological properties of neurons in the LC.

**HIGHLIGHTS:**

- BALB/cJ mice exposed to early life stress are hyperactive in adolescence and displayed attention deficits in adulthood.
- Only female BALB/cJ mice develop increased anxiety in adolescence after early life stress.
- Early life stress causes reduced excitability of locus coeruleus neurons in BALB/cJ mice.
- Only locus coeruleus neurons from female BALB/cJ mice exposed to early life stress displayed an increase in action potential delay time upon depolarization.

## 1. INTRODUCTION

An estimated 44.6% of childhood onset and ∼30% of late onset psychiatric disorders are associated with childhood adversity (Green et al., 2010; McLaughlin et al., 2012). Complex interactions between several factors, including genetic background, sex, and adverse childhood experiences (ACEs) affect susceptibility for a range of psychiatric disorders, including attention deficit hyperactivity disorder (ADHD), anxiety, and depression (Golm et al., 2021, 2020; Rosen et al., 2019; Tibu et al., 2016; Wade et al., 2022). Not only is disruption of monoamine signaling-associated genes implicated in cognitive and affective disorder susceptibility, particularly anxiety disorders, but biological sex and adverse childhood experiences also interact to affect related outcomes (Baumann et al., 2013; Castro Gonçalves et al., 2022; DeYoung et al., 2011; Enoch et al., 2003; Tadic et al., 2003; Voltas et al., 2015). Unfortunately, how genetic predisposition, critically-timed stress, and sex interact to affect susceptibility of developing cognitive and affective processing deficits is unclear.

Based on documented stress-induced change in activity of noradrenergic neurons in the locus coeruleus (LC) (Fan et al., 2014; Keshavarzy et al., 2015; Sheppard et al., 2024) that partly varies based on sex (Bangasser et al., 2013, 2010; Borodovitsyna et al., 2022; Curtis et al., 2006; Jacobson, 2019; Nakamoto et al., 2017; Nishinaka et al., 2015) and the role of these neurons in diverse cognitive and affective processes (Campese et al., 2017; Carlson et al., 2021; Lim et al., 2010; Mair et al., 2005; McCall et al., 2017a; Uematsu et al., 2017) we hypothesize that critically-timed stress exposure may alter processing in this region to shape multiple behaviors. Norepinephrine (NE) released from widely-projecting LC neurons in the central nervous system is well-established as a major modulator of arousal. Decades of work in animal models has indicated that LC neuron projections to the PFC are involved in maintaining attention and arousal (Ishida et al., 2000; Lim et al., 2010; Mair et al., 2005; Pasquier and Reinoso-Suarez, 1978) while those to the basolateral and central amygdala (BLA and CeA) are involved in anxiety and emotional learning behaviors (Campese et al., 2017; McCall et al., 2017b; Uematsu et al., 2017). In mice performing a 5-choice serial reaction time task (5CSRTT), a test to assess attentional performance in animals, inhibiting LC neuron activity using Gi-coupled designer receptors exclusively activated by designer drugs (DREADDs) reduced attentional performance by increasing errors and omissions while inhibition of ventral tegmental area dopamine-containing neurons only affected motivation and response speed (Fitzpatrick et al., 2019). Stimulating LC-NE release in the basal lateral amygdala (BLA) also increases anxiety-like behaviors, indicating a positive feedback loop from the LC to the amygdala that perpetuates anxiety-like behavioral outcomes from stress-associated central inputs (McCall et al., 2017a).

Since data from animal models support a causal role of LC-NE signaling in attention and anxiety and both capacities are also often affected in many disorders of cognitive and affective processing (Jensen and Steinhausen, 2015; Martínez et al., 2016; Saccaro et al., 2021; Shaw et al., 2014; Willcutt et al., 2005) thereby providing a basis for stress-induced disruption of LC-NE function to be involved. Both acute and chronic stress have been shown to alter LC metabolism and NE levels in its projecting brain regions (Fan et al., 2014; Keshavarzy et al., 2015). While multiple synaptic and signaling factors regulate LC neuron activity, levels of corticotropin releasing factor (CRF), the primary neuro-stress signaling molecule, act directly on LC neuron CRF receptor 1 (CRF_1_) to increase LC neuron firing and dose-dependently alter performance in attention set shifting (Cole et al., 2016; Curtis et al., 2006; Snyder et al., 2012). It has been shown that CRF positive inputs from the amygdala increase LC tonic firing to increase anxiety-like and aversive behaviors (McCall et al., 2015). These studies reflect the impact of stress via CRF signaling within the LC on cognitive and affective behaviors. We intend to assess the behavioral changes that occur after our animal model of chronic early life stress and how their behavioral phenotypes relate to LC physiology changes. By determining if there is a correlative relationship between LC physiology with affective/cognitive behavioral deficits, the LC can be further explored as a target for the treatment of ACE-related disorders based on sex and genetic predisposition for anxiety-related disorders.

Given the association of socioemotional stress, especially early in life, with the development of cognitive and affective disorders (Golm et al., 2021, 2020; Green et al., 2010; McLaughlin et al., 2019; Roy et al., 2016; Wade et al., 2022), we hypothesize that stress at critical times produces lasting changes in specific brain regions or circuits that shape cognitive and affective processing, like the LC. In parallel and based on sex-dependent variation in cognitive and affective disorder outcomes (Altemus et al., 2014; Gershon, 2002; Morken et al., 2023), we also expect that these changes will also depend on biological sex. The application of valid pre-clinical animal models to identify mechanisms at play in these interactions is limited, despite the need to understand these mechanisms in order to establish etiologies, prevention, and treatment for a range of cognitive and affective disorders. To address this hypothesis and this outstanding issue of relevant animal models incorporating genetic-determinants of cognitive/affective disruption susceptibility, we sought to develop a model of early life stress-induced behavioral change in an inbred genetic background associated with specific behavioral phenotypes in mice. BALB/cJ mice express increased anxiety-like behaviors relative to C57bl/6J mice (Sartori et al., 2011), increased sensitivity to stress-induced learned helplessness (Jacobson and Cryan, 2007), altered central NE levels after foot shock (Shanks et al., 1994), and contain two polymorphisms within the gene encoding vesicular monoamine transporter 2 (VMAT2) (Crowley et al., 2005). Therefore, we assessed the effect of a novel early life variable stress (ELVS) exposure paradigm (**Figure 1**) on the behavior of male and female BALB/cJ mice as a relevant strain to assess the lasting affect of early life stress in animals with elevated basal anxiety-like behavior. In order to identify potential LC-NE mechanistic changes that may contribute to broad behavioral change, we used electrophysiology to further assessed the synaptic activity and excitability of putative LC neurons in control and ELVS-exposed mice following behavioral testing.

**Figure 1.**
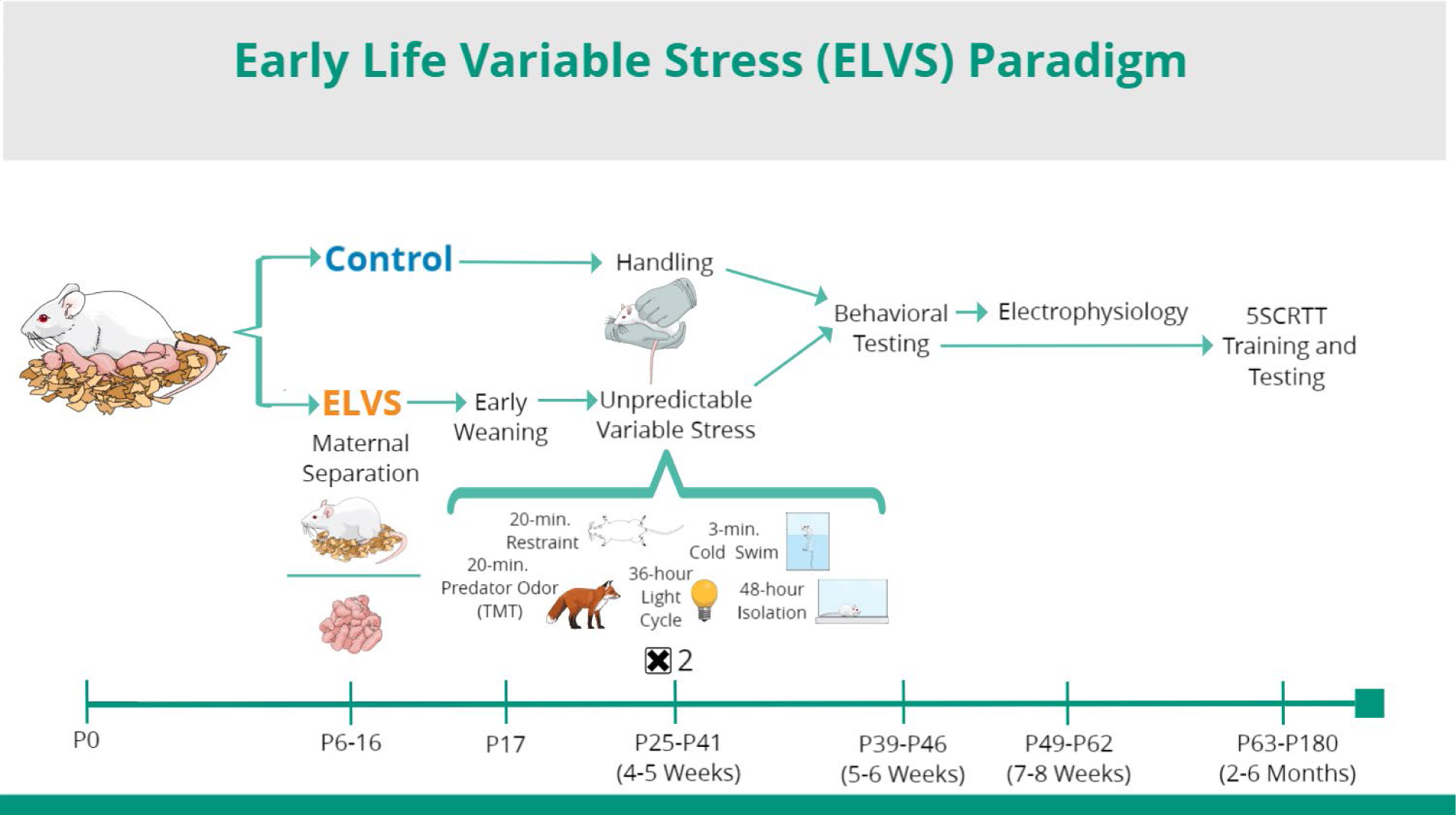
Early Life Variable Stress (ELVS) Paradigm and Experimental Design.

## 2. MATERIALS AND METHODS

### 2.1 Animals

Male and female BALB/cJ (Jackson Labs, strain # 000651) originally purchased from Jackson Laboratory were bred in house. All mice were group housed under a reversed 12-hour light-dark cycle (lights on from 21:00 to 9:00) with *ad libitum* access to food and water. All manipulations and tests were performed during the animals’ active dark phase (9:00 to 21:00), unless otherwise stated. All procedures involving animals were conducted in accordance with protocols approved by the Institutional Animal Care and Use Committee (#2022-100) of Southern Illinois University - School of Medicine.

#### Early Life Variable Stress (ELVS) Animal Model

Male and female BALB/cJ mice were subject to either a two-phase (postnatal and adolescent) early life variable stress (ELVS) paradigm (**Figure 1**) or treated as controls with all mice in a litter assigned to the same group. The ELVS paradigm consisted of two phases of manipulations, first from postnatal day 6-17 and the second from postnatal day 28-40. Mice in the ELVS group were isolated from dams for 4hrs/day during postnatal day 6-10 (P6-10) and weaned early at postnatal day 17 (P17). Pups were supplemented with nutrient gel (Nutra-gel from Bio-serve) from P17 to P21. From P28-42, ELVS mice were subjected to ten consecutive days of stressors with one stress exposure per day, the order of which varied randomly between cohorts/litters. These stressors included 20 minutes of tape restraint (2x), 20 minutes of 2,3,5-Trimethyl-3-thiazoline synthetic predator odor exposure (2x), 3 minutes of forced ice-cold swim in a 40 cm x 10 cm filled container (2x), 36 hours of light cycle disruption via continuous light exposure (1x), and 48 hours of social isolation (1x). Control male and female BALB/cJ mice were not separated from dams prior to weaning, weaned at P21, and were briefly handled 5 times per week for 10-15 minutes beginning at P28.

### 2.2 Innate Behavioral Tests

Behavioral tests were performed in low light conditions (15-20 lux red-light). Mice were habituated to the testing room for 30 minutes prior to performing any assays, which were recorded digitally with a system for video tracking. Each apparatus was cleaned thoroughly with scent-free disinfectant between animals. All data behavioral data were either analyzed by software (EthoVision XT 17.5, Noldus) automatically or by an experimenter blinded to the group of the animal.

#### 2.2.1 Open Field

Mice were placed in a white Plexiglas chamber (40 x 40 cm square open top chamber that is 30 cm high) for 30 min with total distance, time in center (20 x 20 cm region equidistant from the box edges), center entrances, and freezing time calculated based on the center body point of the mouse.

#### 2.2.2 Elevated Zero Maze

Mice were placed in the open arm of an elevated circular platform 5 cm wide and 60 cm off the ground, with 2 parallel walled sections and 2 open sections in between, having an inner diameter of 40 cm. The amount of time spent (mouse center body point) in the open versus closed/walled arms/segments along with open arm entrances were evaluated. Mice entering/spending less time in the open arm were considered to have higher anxiety levels.

### 2.3 Operant Five-Choice Serial Reaction Time Task (5CSRTT)

For 5CSRTT testing of attention capacity, mice were trained and evaluated per standard protocols described here briefly (Fitzpatrick et al., 2018a, 2018b). Two weeks prior to experimentation, mice were water restricted starting at 4 hours of ad libitum access per day which was reduced to 2 hours per day on testing days and ad libitum access on days when testing was not performed. During training sessions, mice were given sweetened condensed milk (800 ms pump time, 20 µL) as a reward for each trial when correctly performing the task and weighed to make sure they were at a proper weight after water restriction. The operant conditioning chamber (30.5 x 24.1 x 29.2 cm) consisting of 2 Plexiglas sidewalls, an aluminum front wall and back wall, and a stainless-steel grid floor was used as the 5CSRTT apparatus (Lafayette Instruments). The front wall contains five nose poke apertures (2.5 x 2.2 x 2.2 cm each) with each aperture containing a light-emitting diode (LED) and an infrared sensor cable to detect mouse nose insertions. The mice were trained (progressive shorting of intertrial interval and stimulus duration) for 30 minutes daily 5-7 days a week to reach baseline performance with intertrial interval (ITI) of 5 seconds and the stimulus duration of 0.8 seconds. In order to pass baseline, mice were required to have an accuracy above 80% and omissions below 30% for three consecutive sessions. After reaching these threshold performance criteria, they were assessed again on three different test variations with either shorter ITI, longer ITI, or shorter stimulus. Response accuracy (% correct) and omitted trials (% omission) for each variable condition was calculated as the primary performance variables.

### 2.4 Ex vivo Acute Brian Slice Preparation

Animals were anesthetized with 4% isoflurane, followed by intracardial perfusion with ice-cold oxygenated N-methyl-d-glucamine (NMDG) artificial cerebrospinal fluid (NMDG-ACSF), which contained (in mM): 92 NMDG, 2.5 KCl, 0.5 CaCl2, 10 MgCl2, 1.2 NaH2PO4, 30 NaHCO3, 20 HEPES, 25 d-glucose, 2 ascorbic acid, 2 thiourea, and 3 sodium pyruvate and had an osmolarity of 300–310 mOsm with pH adjusted to 7.3–7.4 with HCl. Coronal slices (250 μm) that included the LC were acquired in ice-cold NMDG-ACSF with a Compresstome vibrating microtome (Processionary Instruments, LLC) and were transferred to a holding chamber containing NMDG-ACSF at 35 °C where the NaCl concentration increased steadily over 25 minutes. After 30 minutes, slices were transferred to a modified HEPES-based ACSF (HEPES-ACSF, 35 °C), which contained (in mM): 92 NaCl, 2.5 KCl, 2 CaCl2, 2 MgCl2, 1.2 NaH2PO4, 30 NaHCO3, 20 HEPES, 25 d-glucose, 2 ascorbic acid, 2 thiourea, 3 sodium pyruvate, and 3 myo-inositol and had an osmolarity of 300–310 mOsm with pH 7.3–7.4. After incubating slices in HEPES-ACSF at 35 °C for 1 hour, slices were maintained in HEPES-ACSF at room temperature until being transferred to the recording chamber where they were continuously perfused at a 3-5 ml/minute with oxygenated ACSF, which contained (in mM): 125 NaCl, 2.5 KCl, 2 CaCl2, 2 MgCl2, 1NaH2PO4, 26 NaHCO3, 20 D-glucose, 2 ascorbic acid, and 3 myo-inositol and had an osmolarity of 310–320 mOsm with pH 7.3–7.4.

### 2.5 Ex vivo Acute Brain Slice Whole-Cell Electrophysiology

Neurons in the LC were visualized using an upright microscope (Olympus BX51WI) with a 40X water immersive objective. Whole-cell patch-clamp recordings were performed at 32-34 °C maintained with an in-line solution heater from visually identified (infrared differential interference contrast) LC neurons within the LC identified based on regional landmarks and relationship to the 4^th^ ventricle. Using a horizontal puller (P1000, Sutter Instruments), patch pipettes were pulled from filamented borosilicate glass capillaries (outer diameter 1.5 mm, inner diameter 0.86, Sutter Instruments), having a tip resistance of 3–5 MΩ when filled with potassium gluconate-based internal solution that contained (in mM): 139.6 mM K-gluconate, 0.4 mM KCl, 4 mM NaCl, 0.5 mM CaCl_2_, 10 mM Hepes, 5 mM EGTA free acid, 4 mM ATP Mg salt, 0.5 mM GTP Na salt with an osmolality 285–290 mOsm and the pH adjusted to 7.2-7.3 with KOH. For voltage-clamp recordings, 5 mM QX-314 iso-osmotically replaced NaCl in order to block voltage-gated Na+ channels. All signals were acquired at 20 kHz and low-pass filtered at 10 kHz by a Digidata1440 digitizer and a MultiClamp 700B amplifier. Data were collected from the neurons with an input resistance >100 MΩ. If the series resistance for a given recording was >35 MΩ or changed by more than 20% over the course of a recording, the data were rejected for analysis. Liquid junction potentials remained uncorrected in all cases.

### 2.6 Quantification and statistical analysis

Mice of both sexes from control and stressed (ELVS) groups were evaluated for behavioral and electrophysiology experiments. All results are expressed as the mean ± SEM and alpha levels of *p* < 0.05 were considered significant. Two-way ANOVAs (sex, stress) or repeated measures ANOVAs (5CSRTT test or current injection steps, sex, stress) followed by t-tests were used to perform pairwise comparisons between control and ELVS groups within each sex. When data variance was not homogeneous, either a modified t-test was used for pairwise comparisons, a non-parametric Kruskal Wallis H test with a Dunn’s posthoc test was performed in place of two-way ANOVAs, or the alpha level threshold was increased to *p* <0.01 for repeated measures ANOVAs. Statistical test results are not provided for ANOVA main effects or interactions or pairwise comparisons (t-tests) when *p* < 0.1. SPSS 29 (IBM) and Igor Pro 8 (Wavemetrics) were used for automated and manual determination of dependent variable values in EthoVision XT 17.5. Data analyses and graphing were conducted in Clampfit 10.0, Easy Electrophysiology, Igor Pro 8 and SPSS 29.0 (IBM).

## 3. RESULTS

### 3.1 ELVS exposure leads to hyperactivity and female-specific increased anxiety in adolescence

In order to determine how early life stress impacts BALB/CJ mice which display an elevated basal level of anxiety-like behavior, we subjected male and female BALB/cJ mice to an early life variable stress (ELVS) paradigm described above (**Figure 1**). At 6 weeks of age, control and ELVS male and female mice were evaluated on open field (OF) and elevated zero maze (EZM) behavioral assays to assess locomotive and anxiety-like behaviors. The total distance moved in the OF was increased in ELVS-exposed mice regardless of sex. ELVS (male and female) mice moved more than control mice during the 30 minutes of OF testing (**Figure 2A**), suggesting elevated basal activity and exploratory behavior. Although freezing behavior or the number of entries into the center of the OF (**Figure 2B-D**) were not different between groups, there was a strong trend (*p* = 0.055) toward group differences in the amount of time spent in the center of the OF. This trend was driven by a significant reduction in the OF center time for ELVS female mice relative to controls, but not males (**Figure 2E**). In an additional assessment of anxiety, the EZM assay, neither ELVS nor sex affected entries into open arms (**Figure 2F**). However, ELVS exposure significantly affected EZM open arm time regardless of sex (**Figure 2G, H**). Given the sex-specific effect of the anxiety-like behavior measure in the open field and our hypothesis regarding sex-specific changes in behavior due to stress, additional comparisons of stress effects indicated a significant reduction in EZM open arm time (increased anxiety-like behavior) in females, but not males (**Figure 2F-H**). Together, behavior in OF and EZM indicate that ELVS leads to in overall hyperactivity in BALB/cJ mice regardless of sex, but only increases basal anxiety-like behavior in females.

**Figure 2.**
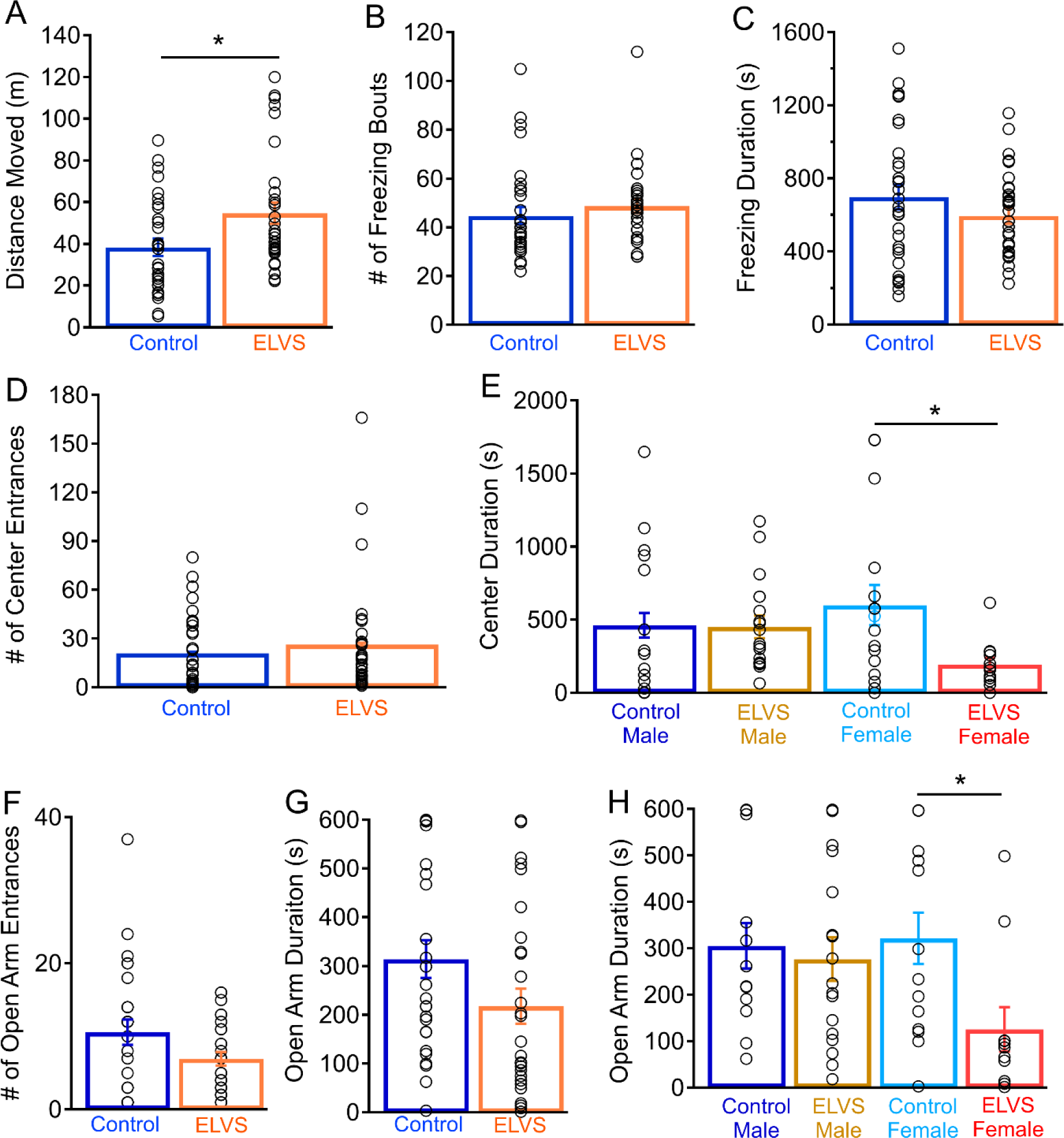
ELVS mice are hyperactive while female ELVS mice have increased anxiety-like behaviors. (**A-E**) Mean ± SEM (bars) and individual (circles) open field (OF) measures for total distance moved in meters (m) (stress: *F*(1,61) = 5.89, *p* = 0.018 *t*(61) = -2.50, *p* = 0.015) **(A)**, number of freezing (**B**), the freezing time (**C**), OF center area entrances for both sexes combined (**D**), and OF center area time (χ^2^(3) = 7.61, *p* =0.055; males: *p* = 0.284; females: *p* = 0.015; **E**). (**F-H**) Mean ± SEM (bars) and individual (circles) values in the EZM, with the number of times mice entered the open arm (stress: *F*(1,53) = 3.57, *p* = 0.065) (**F**), the cumulative time spent in the open arm for data from both sexes combined (stress: *F*(1,53) = 4.47, *p* = 0.039, *t*(52) = 1.81, *p* = 0.077; sex-stress interaction: *F*(d1,53) = 2.49, *p* = 0.12) (**G**), and the time mice spent in the open arm with separated by sex (males: *t*(27) = 0.39, *p* = 0.70; females: *t*(23) = 2.64, *p* = 0.015) (**H**). OF N = 12-19 mice/group, EZM N = 11-17 mice/group, \**p* < 0.05 with pairwise t-tests and two-way ANOVA.

### 3.2 ELVS exposure impairs attention capacity in adulthood

Upon completion of innate behavioral assays, we initiated water restriction and training of a subset of BALB/cJ control and ELVS mice to perform the five serial choice reaction time task (5SCRTT) as an assessment of attention capacity. Stress exposure did not impact the time taken to reach the accuracy and omission threshold criteria during the training period (**Figure 3A**), but males (59.7 ± 4.6 days) took significantly longer than females (43.1 ± 3.8 days) to reach the learning performance criteria as a whole (**Figure 3A**). Measures of baseline performance were taken from the third successful completion of the final training phase (5 second ITI and 0.8 second SD; **Figure 3B & C**), followed by daily testing with three different variable changes with either increased ITI (test 1), decreased ITI (test 2), or shortened stimulus duration (test 3). Although there were no significant accuracy changes due to interactions of testing sequence with sex or sex and ELVS exposure together, there was a significant interaction of ELVS exposure alone with test sequence. Specifically, ELVS exposure led to reduced trial accuracy with increased intertrial interval or decreased stimulus duration (**Figure 3B**). These reductions in performance accuracy on attention tasks that require prolonged maintenance of attention to stimuli (increased ITI and shortened stimulus duration) suggest that vigilance may be perturbed due to ELVS exposure in BALB/cJ mice generally, in addition to hyperactivity.

**Figure 3.**
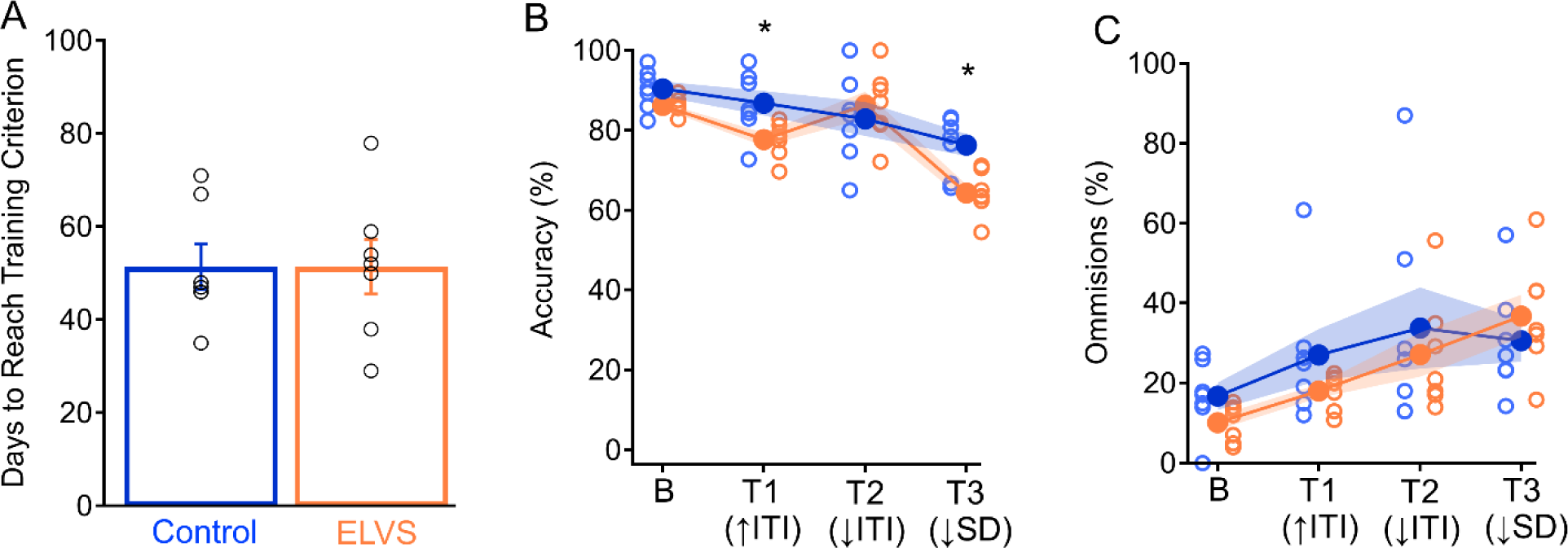
ELVS results in impaired attention in adulthood. **(A)** Mean ± SEM (bars) and individual (circles) number of five choice serial reaction time task (5SCRTT) training days to reach training criterion (sex: *F*(1,13) = 6.722, *p* = 0.027, *t*(12) = 2.79, *p* = 0.016). (**B** & **C**) Mean ± SEM (filled circles with shaded region) and individual (open circles) accuracy (test sequence and stress: *F* (1,13) = 3.74, *p* = 0.022); T1, stress: *t*(12) = 2.60, *p* = 0.023; T3, stress: *t*(12) = 3.43, *p* = 0.005; **B**) and omission percentages (test sequence, stress and sex interaction: *F* (1,13) = 6.49, *p* = 0.002; **C**) on 5CSRTT baseline (B), test 1 (T1) with increased inter trial interval values (↑ITI), test 2 (T2) with reduced inter trial interval values (↓ITI), and test 3 (T3) with reduced stimulus duration values (↓SD). N=7 per group. \**p* < 0.05 with pairwise t-tests and repeated measures three-way ANOVA.

### 3.3 ELVS alters spontaneous postsynaptic inhibitory currents in LC neurons of male mice

Given the functional reciprocal connectivity of the LC with the amygdala and PFC to shape both anxiety-like and attentional behaviors, we hypothesized that changes in central adrenergic system may underpin these altered behaviors. First, to assess spontaneous excitatory and inhibitory synaptic activity within LC neurons, we performed voltage clamp electrophysiology in putative LC neurons from acute brain slices *ex vivo*. In order to isolate spontaneous excitatory post synaptic currents (sEPSCs), neurons were voltage-clamped at -60 mV which was equal to the E_Cl_ (**Figure 4A**). We did not observe any significant differences in sEPSC amplitude or frequency (**Figure 4B & C**). To next isolate spontaneous inhibitory postsynaptic currents (sIPSCs), putative LC neurons were voltage-clamped at 0 mV which was near the cation reversal potential (**Figure 4D**). Evaluation of sIPSCs identified a significant increase in sIPSC frequency in ELVS male mice relative to male controls without stress-related changes in sIPSC activity in females (**Figure 4E & F**).

**Figure 4.**
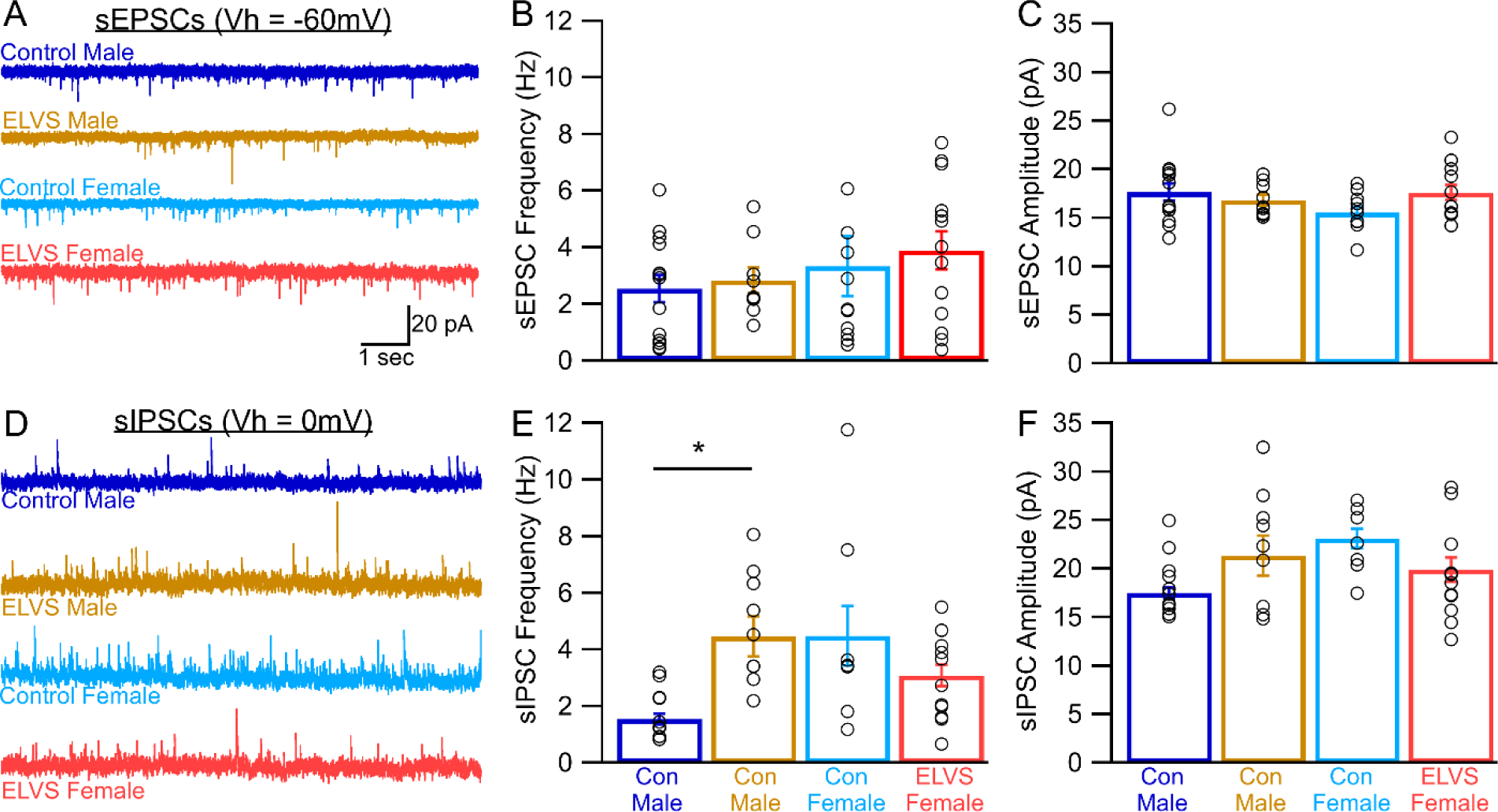
ELVS selectively increases inhibitory synaptic tone in male LC neurons. (**A, D**) Representative voltage clamp recording traces of control and ELVS putative LC neuron spontaneous excitatory postsynaptic currents (sEPSCs, Vhold = -60mV, **A**) and inhibitory postsynaptic currents (sIPSCs, Vhold = 0mV, **D**). (**B, C**) Mean ± SEM (bars) and individual (circles) sEPSC frequency (**B**) and amplitude (**C**). (**E, F**) Mean ± SEM (bars) and individual (circles) sIPSC frequency (**E**) and amplitude (**F**) indicate sIPSC frequency (sex-stress: *F* (1,35) = 8.25, *p* = 0.007) is selectively elevated in male ELVS (*t*(16) = 3.80, *p* = 0.002) LC neurons with sIPSC amplitude (sex-stress interaction: *F* (1,35) = 5.34, *p* = 0.027) not different between groups (t-test *p* > 0.05). N = 3 mice per group, sEPSC: n = 9-13 cells/group, sIPSC: n = 8-10 cells/group, \**p* < 0.05 with pairwise t-tests and two-way ANOVA.

### 3.4 ELVS LC neurons display reduced excitability and input resistance

We next evaluated LC neuron excitability and additional membrane properties. ELVS mice displayed reduced current injection-evoked excitability in both sexes (**Figure 5A & B**). Both stress exposure and sex affected spontaneous firing rate of LC neurons, which was reduced in both male and female ELVS mice relative to control mice (**Figure 5C & D**). We also observed a sex-specific changes in the resting membrane potential, with ELVS causing membrane potential hyperpolarization males and depolarization in females (**Figure 5E**). Without significant changes in the input/membrane resistance (**Figure 5F**) of LC neurons, this resting potential change in male ELVS mice relative to controls accounts for evoked and spontaneous firing rate changes observed in this group. However, these changes alone, without meaningful changes in synaptic drive in LC neurons from female mice exposed to ELVS failed to explain the reduced excitability observed in this group.

**Figure 5.**
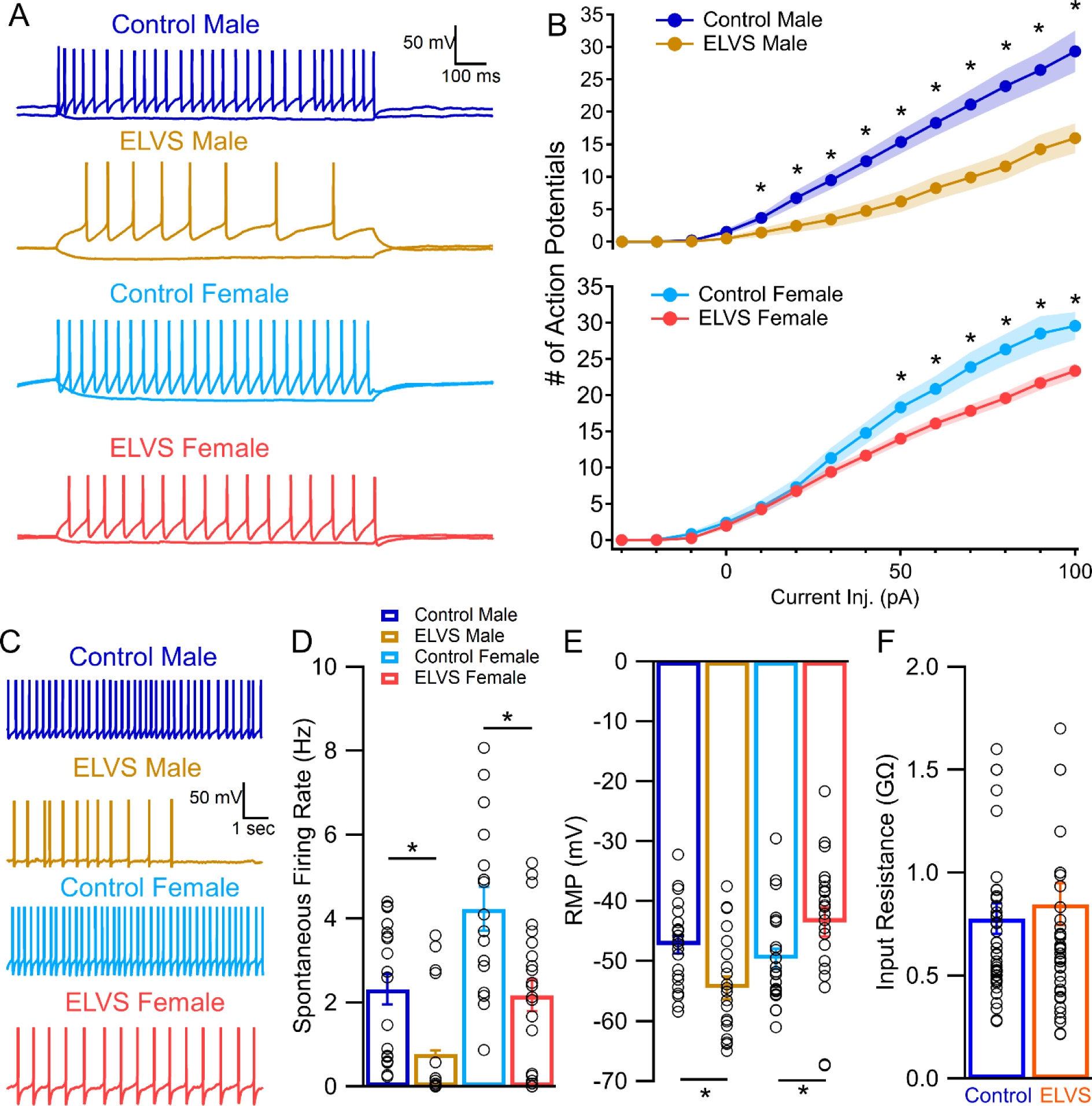
ELVS results in reduced LC neuron excitability input resistance. (**A**) Representative traces of current clamp recordings from control and ELVS putative LC neurons in response to +80/-30 pA current injection (1 s) from each neuron’s natural resting membrane potential. (**B**) Mean ± SEM (filled circles with shaded region) action potential (AP) number in response to current (pA) injection at each level for males (upper) and females (lower) of each group (sex: *F*(1,83) = 8.41, *p* = 0.005; stress: *F*(1,83) = 19.22, *p* < 0.001). (**C**) Representative traces of spontaneous firing (10 s) recordings from putative LC neurons from each group. (**D-F**) Mean ± SEM (bars) and individual (circles) spontaneous firing rates (sex: *F*(1,72) = 14.50, *p* < 0.001; stress: *F*(1,72) = 17.29, *p* < 0.001; males: *t*(33) = 2.62, *p* = 0.013; females: *t*(36) = 3.29, *p* = 0.002; **D**), resting membrane potential (sex: *F*(1,86) = 4.65, *p* = 0.034; sex-stress interaction: *F*(1,86) = 1170, *p* = 0.001; males: *t*(41) = 2.82, *p* = 0.007; females: *t*(42) = - 2.14, *p* = 0.038; **E**), and input/membrane resistance (**F**). n = 14-24 cells/group from N = 3 mice/group. \**p* < 0.05 with pairwise t-tests and three-way repeated measures ANOVA (**B**) or two-way ANOVA (**D-F**).

### 3.5 ELVS LC neurons show a sex-specific increase in action potential delay time

Since resting potential changes with ELVS, likely due to increase inhibitory drive (**Figure 4**), only explained excitability reductions in LC neurons of male mice, we next assessed LC neuron evoked excitability when the resting membrane potential was adjusted with constant current injection to -60 mV (∼E_Cl_, to nulify the effects of inhibitory drive; **Figure 6A**). With an initial resting potential of -60 mV, there remained a significant effect of ELVS exposure on current injection-evoked firing across the range of current values in LC neurons from both sexes (**Figure 6B**), but the interaction between sex and ELVS exposure observed at natural resting potentials was lost. However, for consistency, the data are shown with each sex separate where it becomes clear that stress effects in male are comparable at natural resting potential or at -60 mV, but this configuration identifies greater changes in excitability due to ELVS exposure in females (**Figure 6B**). It also became clear that, unlike at more depolarized natural resting potentials, putative LC neurons specifically from ELVS female mice displayed a delay before the first action potential across nearly all current injections (**Figure 6A**). A comparison of first action potential delay times across groups for all neurons eliciting action potentials in response to 70-100 pA injection identified a sex and ELVS exposure interaction exposure with the delay time being significantly different between control and ELVS in LC neurons form female mice in response to each current injection, but not males (**Figure 6C**).

**Figure 6.**
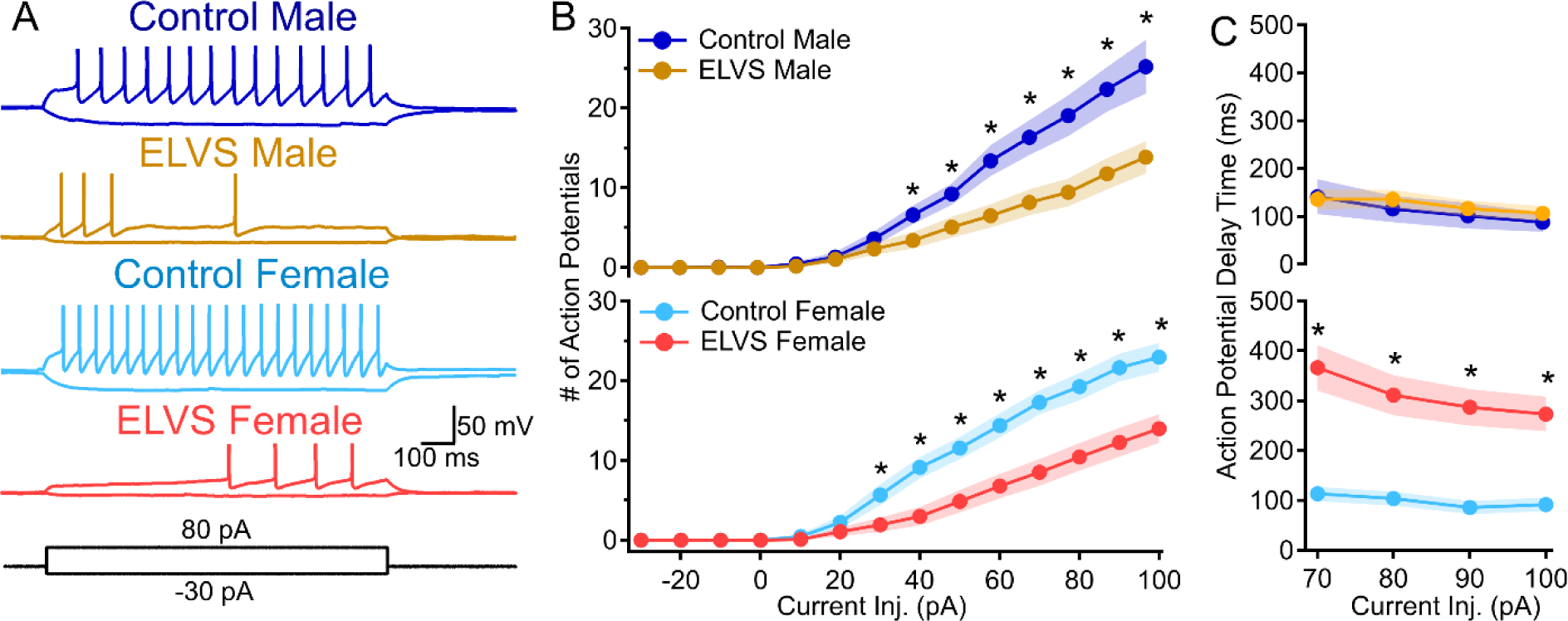
ELVS results in a female-specific increase in LC neuron action potential delay time. (**A**) Representative traces of current clamp recordings from control and ELVS putative LC neurons in response to +80/-30 pA current injection (1 s) with the initial membrane potential adjusted to -60 mV with current injection. (**B-C**) Mean ± SEM action potential (AP) number (stress: *F*(1,74) = 18.49, *p* < 0.001) (**B**) and first action potential delay time (**C**) due to 70-100 pA current injection with a starting membrane potential of -60 mV (sex-stress interaction: *F*(1,69) = 13.90, *p* < 0.001). n = 14-23 cells/group, N = 3 mice/group, \**p* < 0.05 with pairwise t-tests and three-way repeated measures ANOVA.

## 4. DISCUSSION

### 4.1 Summary of Findings

To develop an animal model in order to understand the underlying mechanisms involved in early life stress-induced changes in affective and cognitive behaviors, we found that combining deprivation in early life and unpredictable stress in adolescence resulted in hyperactivity along with female-specific increased anxiety-like behavior in adolescence and impaired attention in both sexes persisting into early adulthood. Paralleling behavioral deficits, LC neuron excitability was reduced in mice of both sexes after early life stress through different mechanisms. While only LC neurons in female ELVS mice displayed a dramatic increase in action potential delay time likely due to differential activation of voltage-gated ion channels, LC neurons of male mice exposed to ELVS were more hyperpolarized. These findings suggest that early life stress impacts LC physiology in sex-specific ways that may contribute to persistent attention deficits and/or anxiety-like behavior.

### 4.2 The LC as an interface between stress and affective/cognitive behaviors

The LC is innervated by CRF-containing neuron terminals from the paraventricular hypothalamic nucleus (PVN), the CeA, the Barrington’s nucleus, and the nucleus paragigantocellularis (Koegler-Muly et al., 1993; Valentino et al., 1992). Intracerebroventricular CRF injection into the brain and direct CRF application onto LC neurons has been shown to increase LC neuron spontaneous firing rates (Valentino et al., 1983) via activation of CRF_1_ receptors (Sánchez et al., 1999). CRF_1_ couples to G_i_, G_s_, and G_q_ pathways, with notable sex differences in preferred coupling and receptor internalization shown in the LC in rodent studies (Bangasser and Wiersielis, 2018; Curtis et al., 2006). It is perhaps these sex-differences in CRF_1_ singling in the LC that may lead to sex-specific persistent changes in basal LC physiology affecting inhibition and excitability identified here. The LC is involved in both anxiety and attention, primarily due to its modulation of the amygdala and PFC (Aston-Jones et al., 1999; Campese et al., 2017; Lim et al., 2010; Mair et al., 2005; McCall et al., 2017a; Van Stegeren et al., 2005; Zhang et al., 2023). The consistent reduction in noradrenergic LC neuron excitability in male and female BALB/cJ mice exposed to ELVS may provide a window into mechanisms by which prolonged stress early in life produces lasting and broad reach effects on neural function. The addition female-specific increase in the action potential delay time when cells were held at a -60 mV further indicates that there may be stress-sensitive mechanistic changes that would alter excitability with or without changes in input (Jerng et al., 2004). Rather the female-specific changes in LC neurons of ELVS mice would largely affect how excitatory input is integrated to modulate LC neuron firing over time. Since it has been shown that LC neurons must transition from different firing modes (i.e. tonic vs. phasic) in order to shift into different optimal attention states depending on the task (Howells et al., 2012), changes in how these transitions occur may rely on mechanisms that shape both spontaneous and evoked excitability.

### 4.3 Conclusions and Future Directions

Exposure to stress, especially at critical periods in early life, has clinically relevant impacts on behavior neuropsychiatric disorder risk. In general, women have a greater risk for stress related disorders. Women are twice as likely to have depression, GAD, and post-traumatic stress disorder (PTSD) when compared to men, with this difference emerging after adolescence (Kessler et al., 2004; Morken et al., 2023). Sex differences in adolescent stress effects on affective behaviors could explain the sex differences in anxiety-like behavioral outcomes seen in our model. Typically, boys more commonly suffer from neurodevelopmental disorders such as ADHD or autism that are also associated with ACEs (Bölte et al., 2023). In childhood, the male to female ratio of ADHD prevalence is 3:1, but approaches 1:1 in adulthood (Bölte et al., 2023; da Silva et al., 2020). In this study, 5SCRTT attention testing is performed during adulthood, suggesting that in both males and females there are lasting attentional deficits in our ELVS model. Further investigation regarding behavioral outcomes when isolating the early life and adolescent components of the ELVS paradigm will help to understand if each component is necessary for behavioral deficits, if a certain component is responsible for sex-specific effects, or if only one component is needed for the behavioral outcomes we have observed.

The results of this study support further exploration into LC-NE related treatment targets for ACE-related disorders, particularly when attention deficits are present. It also suggests sex is a crucial biological variable involved in determining psychiatric outcomes due to ACE(s) exposure and treatment. This study suggests early life stress results in reduced LC neuron excitability in both sexes, but the causal mechanism and functional impacts of this change may differ between males and females resulting in sex differences in behavioral outcomes. Further investigation of whether similar treatment targets will work in both sexes or if more specific targets depending on sex may be more beneficial at ameliorating affective and/or cognitive behavioral deficits.

## DECLARATION OF COMPETING INTEREST

The authors declare that they have no known competing financial interests or personal relationships that could have appeared to influence the work reported in this paper.

## ETHICS APPROVAL

All procedures involving animals were performed in accordance with protocols approved by the Institutional Animal Care and Use Committee at Southern Illinois University – School of Medicine.

## AVAILABILITY OF DATA AND MATERIALS

The datasets used and/or analyzed during the current study are available from the corresponding author on reasonable request with statistical analysis results included with this published article’s supplementary information files.

## FUNDING

This work was supported by a Research Seed Grant and laboratory operating funds provided to BDR by Southern Illinois University – School of Medicine.

## ABBREVIATIONS

ACEs: Adverse Childhood Experiences
ADHD: Attention deficit hyperactivity disorder
GAD: Generalized anxiety disorder
NE: Norepinephrine
LC: Locus coeruleus
CRF: Corticotropin-releasing hormone/factor
ELVS: Early life variable stress
VMAT2: Vesicular monoamine transporter 2
PFC: Prefrontal cortex
5CSRTT: 5-choice serial reaction time task
DREADDs: Designer receptors exclusively activated by designer drugs
CeA: Central amygdala
p-ERK: Phosphorylated extracellular signal- regulated kinase
CRF_1_: CRF receptor 1
OF: Open field
EZM: Elevated zero maze
sEPSCs: Spontaneous excitatory post synaptic currents
sIPSCs: Spontaneous inhibitory postsynaptic currents
PVN: Paraventricular hypothalamic nucleus
PTSD: Post-traumatic stress disorder

## ACKNOWLEDGEMENTS

The authors would like to thank Hunter McKinney and Alan Huebschen for their assistance in collecting behavioral assay data.

## AUTHORS’ CONTRIBUTIONS

Savannah Brannan: Conceptualization, Methodology, Validation, Formal Analysis, Investigation, Data Curation, Writing – Original Draft, Writing – Review & Editing, Visualization, Lauren Garbe: Formal Analysis, Ben D Richardson: Conceptualization, Methodology, Validation, Data Curation, Writing – Review & Editing, Visualization, Supervision, Project Administration, Funding Acquisition.

## Notes

### Competing Interest Statement

The authors have declared no competing interest.

